# Brain-responsive music enables non-invasive, targeted and unobtrusive neurostimulation

**DOI:** 10.1101/2024.07.30.605888

**Authors:** Boubker Zaaimi, Rosie Clay, Abhishek Shivakumar, Joern Rickert, Andrew Jackson

## Abstract

**Objective:** We are developing a new closed-loop brain stimulation method by embedding, within music, auditory elements that respond to the listener’s brain activity. Here we show that this brain-responsive music has systematic and targeted effects on neural oscillations implicated in a variety of neurological and mental health disorders.

**Approach:** We recorded magnetoencephalogram (MEG) or electroencephalogram (EEG) signals from participants as they listened to music synthesized by commercial audio software. Brain signals were bandpass filtered, phase-shifted and used to control the timbre and/or timing of notes within the music.

**Main results:** Listening to brain-responsive music induced peaks and troughs in spectral power at frequencies that depended systematically on the phase-shift applied to the brain signal. Phase-dependent modulation was greatest at the centre frequency of the filter. As a result, by calibrating these parameters we could achieve selective enhancement or suppression of either theta (5 Hz) or alpha (10 Hz) oscillations. Moreover, by chosing different sensor locations we could target power modulation to either frontal or temporal cortex. The phase-dependent power modulation observed with brain-responsive music was significantly attenuated when participants listened to identical music as a conventional, open-loop stimulus. Finally, we demonstrate that brain activity could be modulated by more complex compositions combining a variety of brain-responsive musical elements controlled by a wireless, wearable EEG headband suitable for home use.

**Significance:** Brain-responsive music provides an unobtrusive and targeted method of modulating neural oscillations in the listener’s brain, and may enable both creative and therapeutic applications of Brain Computer Interface technologies.

## 1. Introduction

Many neurological and mental health conditions are characterised by altered network dynamics, manifesting as abnormal oscillatory activity in different brain areas and frequency bands[1]. A promising therapeutic approach is to regulate activity patterns with neurostimulation, either enhancing under-active oscillations or suppressing over-active frequencies. While traditional neurostimulation therapies deliver open-loop stimuli in predefined patterns, for example as fixed-frequency trains, there is increasing interest in closed-loop methods whereby stimulus parameters respond to the user’s brain activity sensed by Brain-Computer Interface (BCI) technologies[2]. From a theoretical perspective, closed-loop feedback should enable more precise regulation of brain dynamics compared to open-loop, feedforward control, since the effect of a stimulus will depend on the instantaneous state of the target network. As a result, closed-loop electrical or optogenetic stimulation delivered at specific phases of ongoing brain rhythms can either enhance or suppress pathological activity in epilepsy[3] and Parkinson’s disease[4]. Similarly, closed-loop auditory stimulation (CLAS) can manipulate slow oscillations in sleep to influence memory consolidation[5], and has recently been applied to alpha activity in awake subjects to modify sleep onset dynamics[6]. Closed-loop therapies have additionally been proposed for conditions including Alzheimer’s disease, schizophrenia, anxiety disorders, depression and addiction[7]. An advantage of the auditory modality over electrical stimulation for closed-loop applications is the absence of stimulus artefacts, simplifying the task of real-time control with non-invasive MEG or EEG. However, to date, CLAS protocols have been limited to simple stimuli such as white noise bursts, with only the timing of each burst controlled by brain signals. In the awake state, such repetitive sounds may be distracting or annoying over extended periods of time, and are likely to have a relatively weak effect on brain areas beyond the auditory system.

Music represents another powerful approach to influencing the brain with auditory stimuli, and is arguably our oldest and most sophisticated neurostimulation technology. Ever since our Neanderthal ancestors developed primitive instruments[8], we have used music to rouse and relax, express joy and sadness, to help us go to war and to fall in love. In contrast to CLAS protocols, music incorporates many different structural elements (rhythm, melody, harmony) and expressive cues (articulation, timbre) that can profoundly influence the listener’s affective state. Whether these associations are innate or acquired, listening to music activates widespread brain areas, including limbic, autonomic and motor networks[9, 10]. As a result, therapeutic applications of music, from pain relief, improved sleep and memory to the treatment of depression and anxiety, are increasingly recognised[9, 11]. Moreover, music exerts both direct and indirect influences on neuronal oscillations. For example, the rhythms in music are thought to entrain the phase of oscillations in auditory areas[12, 13] while the valence and arousal levels of music can affect the power of spectral features including frontal midline theta[14] and alpha asymmetry[15]. However, passive music listening can be considered similar to open-loop brain stimulation in that the pattern of auditory stimuli is entirely determined by the composer and performers rather than responding to the state of the listener’s brain.

CLAS protocols (feedback control with simple, low-dimensional stimuli) and conventional music (open-loop delivery of complex, high-dimensional stimuli) can thus be considered opposing ends of a spectrum (Figure 1a). Our research aims to integrate these approaches by applying the scientific principles of closed-loop stimulation to the rich and expressive textures of music in order to modulate brain oscillations for therapeutic purposes. Our vision of brain-responsive music leverages emerging consumer BCI technologies such as wearable, wireless EEG headsets, together with the audio capabilities of mobile devices, to create music that is personalised to the real-time state of the listener’s brain. In effect, we aim to exploit both the power of music and the precision of closed-loop feedback to deliver a new, versatile and targeted form of neurostimulation.

**Figure 1.**
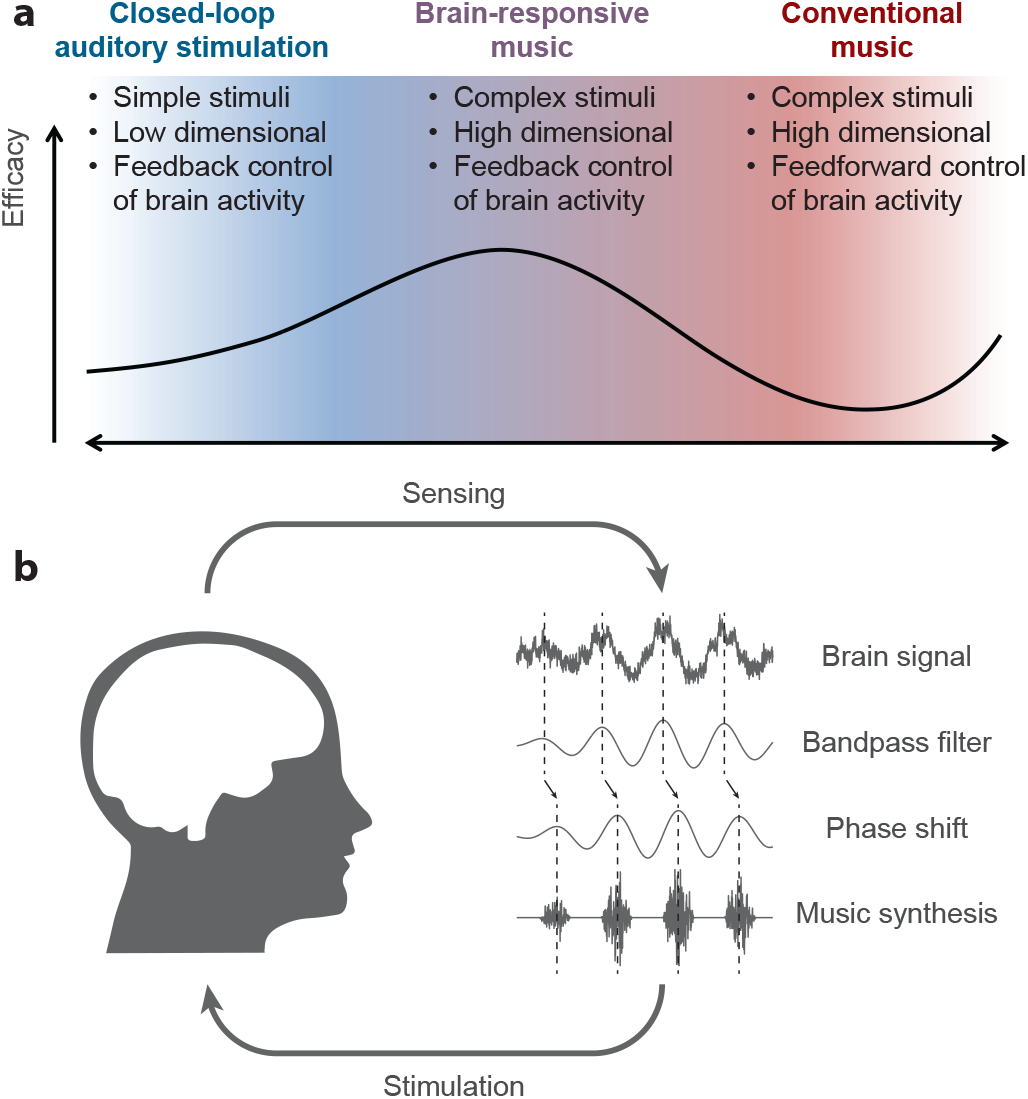
Brain-responsive music concept and schematic. (a) Both closed-loop auditory stimulation and conventional music influence the brain with acoustic stimuli, but sit on opposite ends of a spectrum from feedback control of simple stimuli to feedforward delivery of complex stimuli. We hypothesize that brain-responsive music, which combines feedback control with complex stimuli, could provide greater efficacy for modulating brain activity. (b) Our approach to brain-responsive music establishes a feedback loop between the brain and music. A phase shift imposed within this loop tunes its properties to positive or negative feedback.

Towards this ambition, we have developed methods to connect BCI hardware with music synthesis software capable of generating a variety of real-time brain-responsive music. Our results demonstrate that brain-responsive music can systematically enhance or suppress neural oscillations at targeted frequencies with spatial specificity. Interestingly, these effects are not observed when subjects listen to identical music as a conventional ‘open-loop’ stimulus. Moreover, we show that brain oscillations can be modulated using a simple wearable, wireless headset suitable for home use. By enabling music creators to produce brain-responsive music within familiar digital audio environments, we hope to facilitate a creative collaboration between art and science as well as develop a new non-invasive and non-intrusive neurostimulation modality with the potential for widespread adoption.

## 2. Methods

### 2.1 General overview

Conceptually, our approach creates a feedback control loop using music to deliver auditory stimulation to the brain at specific phases of endogenous neural oscillations. By tuning the properties of this feedback loop, we aim to enhance or suppress the amplitude of ongoing oscillations, much like pushing on a swing at the appropriate moment in its cycle. Figure 1b shows a schematic of the processing steps within this feedback loop, which are described in detail in the sections below.

### 2.2 Hardware setups

For any closed-loop controller, delays in the signal pathway are a critical determinant of the overall performance that can be achieved. Fortunately, we can leverage powerful tools that are commonplace in modern music production workflows and increasingly available on mobile devices. Digital Audio Workstation (DAW) software hosts virtual instruments and audio effects, implemented in standardized architectures such as Virtual Studio Technology (VST3) plugins, that process incoming audio and Musical Instrument Digital Interface (MIDI) data. Since musicians playing live are acutely sensitive to latency, jitter and sample dropout, DAW and plugin software, audio interfaces and drivers are optimised for fast and reliable throughput of data at high sampling rates, making them ideal also for closed-loop brain-responsive music applications. For example, a typical input/output buffer size of 256 samples at 48 ksp/s audio rate entails just 10 ms of roundtrip latency to process incoming data and synthesize complex music comprising multiple instruments and effects.

The experiments reported here used Live 11 Suite (Ableton) software to generate brain-responsive music from MEG or EEG signals. This popular DAW can receive incoming data from three sources: analog input from an audio interface (in our case an M4, MOTU), MIDI data from an external device, or Open Sound Control (OSC) messages sent over a network connection. We exploited these in three different experimental setups (Figure 2a):

**Figure 2.**
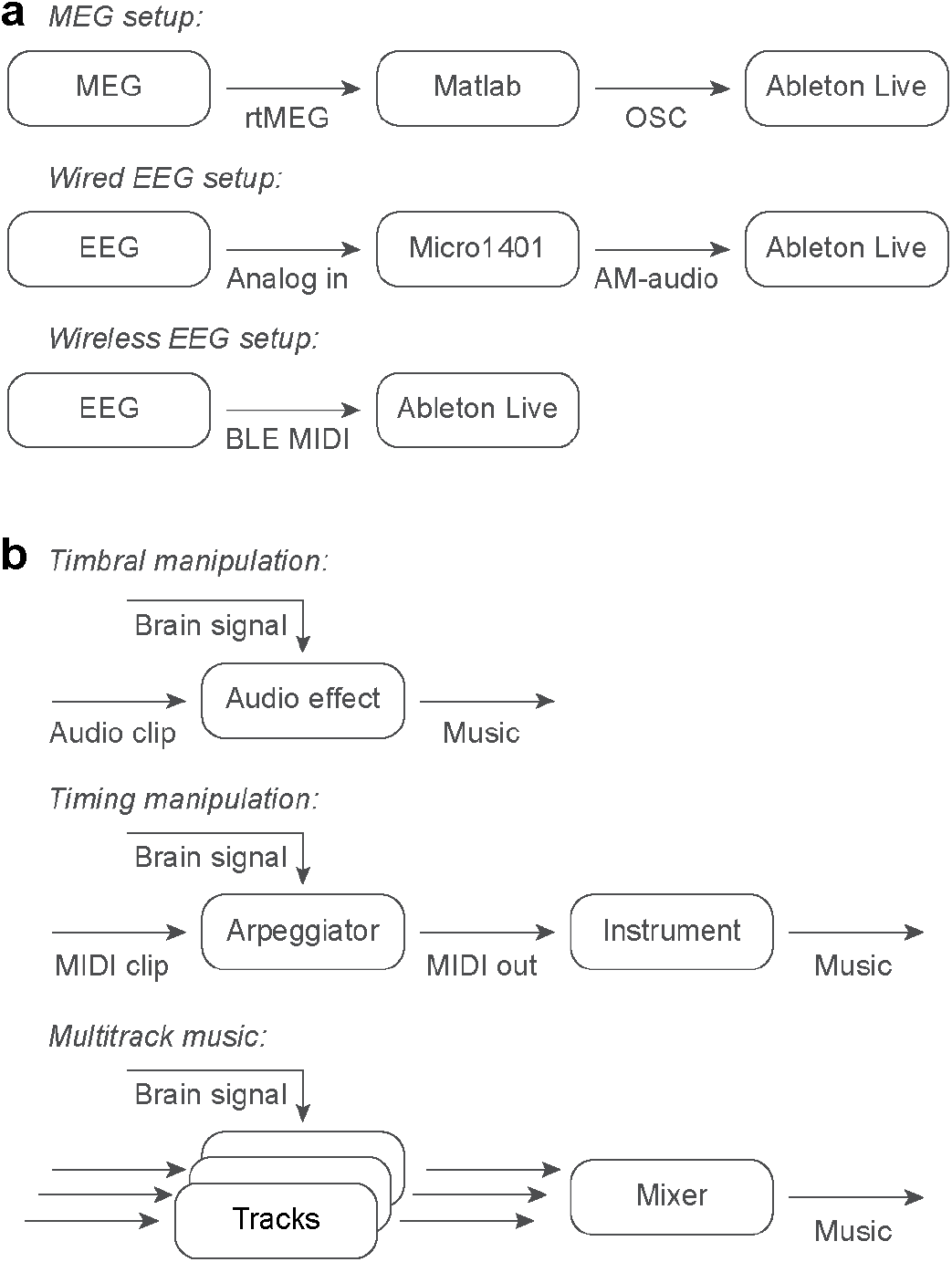
Methods for generating brain-responsive music. (a) Signal pathways enabling brain control of music synthesis by Ableton Live software. (b) Within Ableton Live, the brain signal can be used to modulate the timbre or timing of notes within multitrack compositions.

#### MEG setup

Signals were aquired using a Neuromag (Elekta) MEG system. We used rtMEG software[16] to stream data at 1000 sp/s to a FieldTrip[17] buffer, and custom Matlab (MathWorks) code to convert data from one of 204 gradiometer channels into OSC messages sent to Ableton Live. Subjects listened to the resultant music through MEG-compatible pneumatic earphones (TIP-300, Nicolet Biomedical Instruments).

#### Wired EEG setup

A single channel of differential EEG was recorded between adhesive electrodes (Red Dot 26600-5, 3M) applied to the forehead (approximately AFz) and right mastoid, with a ground electrode on the neck. EEG was amplified (NL824, Digitimer) and sampled at 500 sp/s using a Micro1401 (CED). Because the audio inputs of the M4 audio interface incoporate high-pass filters, we configured the Micro1401 to output the EEG signal amplitude-modulated onto an audio-frequency carrier. Subjects listened to the music on wired, closed-back headphones (SH-450, Focusrite).

#### Wireless EEG setup

We additionally developed a custom headband to amplify and digitise a differential EEG signal and relay it directly to Ableton Live via a Bluetooth Low Energy (BLE) MIDI protocol. Our headband used a single dry electrode (Dryode, IDUN) on the forehead and Ag-AgCl earclips (OpenBCI) on the right/left earlobes for reference/ground. Our wearable electronics incorporated front-end amplification (2xAD623, Analog Devices) and a microprocessor (PIC16F1705, Microchip Technology) to digitise the signal at 333sp/s and convert it into MIDI control change (CC) messages, which were relayed to a BLE transmitter (WIDI Core, CME). For the experiment reported here, the subject listened to the music on the in-built speaker of a MacBook Pro laptop.

### 2.3 Signal processing

To generate a signal to modulate music synthesis from incoming MEG or EEG data we used a filter/phase-shift algorithm that we have previously described for closed-loop optogenetic stimulation[3]. Briefly, a single incoming brain signal is band-pass filtered and phase-shifted using a finite impulse response (FIR) filter with a convolution kernel of the form:

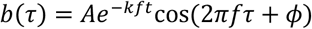

where *f* determines the centre frequency of the band-pass filter (here we used either 5Hz, 6Hz or 10Hz), *k* (in the range 0.8-1.25) determines its width and *ϕ* is the relative phase advance from input to output. This phase shift, in combination with the signal processing time, audio synthesis latencies and physiological delays in the brain’s response to the music, determines the total phase shift in the feedback loop between the brain and music. A closed-loop phase shift of zero will result in positive feedback and enhancement of activity at frequency *f*, while a closed-loop phase shift of 180° will result in negative feedback and suppression of that activity.

For the MEG experiments this algorithm was implemented within the Matlab code that streamed sensor data from the FieldTrip buffer. For the EEG experiments, we developed a custom VST3 plugin using the JUCE C++ framework to implement the algorithm inside Ableton Live. This plugin resampled incoming EEG data (either demodulated audio or MIDI CC messages) to 750 sp/s and applied the FIR filter using a kernel length of 2048 samples (2.7 s).

### 2.4 Brain-responsive music synthesis

Music is a rich and complex stimulus space, and a DAW environment like Ableton Live presents open-ended possibilities for modulating synthesis parameters with brain signals. Responses to music are influenced both by individual tastes and cultural conventions, so determining a single stimulus manipulation that is optimal for all listeners is likely futile. Rather we aim to develop tools to embed brain-responsive elements into a range of musical styles so as to preserve creative flexibility for composers and the aesthetic qualities of music. For the proof-of-concept experiments presented here we tested simple manipulations of both the timbre and timing of musical elements using the inbuilt functionality of Ableton Live (Figure 2b):

#### Timbral manipulations

The timbre of a musical element is determined by its harmonic overtones which can be manipulated by filtering. Our first experiments used a resonant low-pass filter (‘Auto Filter’ device) applied to a simple repeating chord progression. The output of our closed-loop algorithm was halfwave rectified and mapped to the cut-off frequency of the filter to create a brain-responsive ‘wah-wah’ effect (Supp File 1).

#### Timing manipulations

A straightforward way to manipulate the timing of musical elements is to alter MIDI data as it is passed to a virtual instrument. For this we developed a custom ‘arpeggiator’ device using the inbuilt programming language Max4Live. A conventional arpeggiator takes incoming chords and plays each note individually in a rhythmical ascending or descending pattern. Our device similarly takes incoming chords and plays each note individually, but instead timed to the peak of the output of our closed-loop algorithm, with a loudness scaled by the amplitude of that peak. We used a freely-available collection of short MIDI pieces (‘Pretty Piano Progressions’, Caelum Audio), passed via the brain-responsive arpeggiator into a virtual piano instrument (‘Upright Piano by Spitfire Audio’). Each piece comprised a eight bar progression repeated twice at 120bpm to produce 30s clips of brain-responsive music (Supp File 2).

While simple, these manipulations provide building blocks that can be used to construct more sophisticated and interesting multitrack brain-responsive music. To begin exploring these possibilities, we additionally report an extended composition in which the timing and timbre of multiple instruments are manipulated in different ways by the incoming brain signal (Supp File 3).

### 2.4 Experimental protocols

Experiments were performed with volunteers with no history of neurological disorders, under appropriate local ethical approval from Aston University or Newcastle University. Participants listened to brain-responsive music at a comfortable volume with their eyes closed. Epochs of different experimental conditions (between 10-30s in duration) were presented multiple times in pseudorandomised order (between 1-10min of music per condition in total). Across the experiments reported here, these conditions generally comprised different phase-shifts applied within the closed-loop algorithm (e.g. 45°, 90°, …, 360°). We additionally incorporated various control conditions within different experiments:

#### Baseline epochs

For timbral manipulation experiments, we included epochs where the input to the filter/phase-shift algorithm was held at zero, so that the music was played unmodulated. For the timing manipulation experiments, holding the input at zero resulted in silence (since the arpeggiated notes were triggered by peaks in the output signal). Therefore our baseline epochs instead comprised white noise input to the filter/phase-shift algorithm, resulting in a random timing of notes with approximately the same tempo as in closed-loop conditions.

#### Replay epochs

We also wished to generate a control condition that precisely matched the temporal structure of music under closed-loop conditions. Therefore we used brain signals recorded from the same subject during closed-loop epochs, replayed a second time into the filter/phase-shift algorithm. In this way, we generated music with identical manipulations to the closed-loop epochs, but these were no longer synchronized to endogenous brain activity. Since the spectrum of the brain signal (and hence the temporal structure of the resultant music) depended on the closed-loop phase-shift, we included separate replay conditions matched to each of the experimental phase-shift conditions.

MEG experiments comparing different filter frequencies and source channels were performed in separate blocks, in counterbalanced order across subjects.

### 2.4 Analysis methods

Analysis was performed with custom Matlab and Python scripts using Fieldtrip[17] and MNE-Python[18] toolboxes. Source reconstruction was performed using the dSPM algorithm to estimate power spectra for a mesh of 8196 cortical sources co-registered to the ‘fsaverage’ template. Average power spectra in either sensor or source spaces were compiled for each closed-loop condition and expressed as decibels relative to the baseline condition. For each frequency, this relative power was fit with a sinusoidal function of the closed-loop phase-shift to determine the phases associated with maximal enhancement/suppression as well as the peak-to-trough depth of modulation.

To assess the statistical significance of power modulation across the whole head, we pooled all subjects to calculate an overall modulation depth which was compared to a null distribution constructed after shuffling phase conditions for each subject.

## 3. Results

### 3.1 Brain-responsive music can enhance or suppress oscillatory power

Our initial experiments established that music synchronized in real-time to specific phases of ongoing neural oscillations could enhance or suppress spectral power at specific frequencies. Figure 3a shows power spectra for an MEG signal recorded from the right temporal cortex of a representative participant in a timbral manipulation experiment (see Methods). This signal was, in real-time, band-pass filtered at 5 Hz, phase-shifted by one of eight equally spaced angles, and used to modulate an audio filter (‘wah-wah’) effect applied to the music that the subject was listening to (Supp File 1).

**Figure 3.**
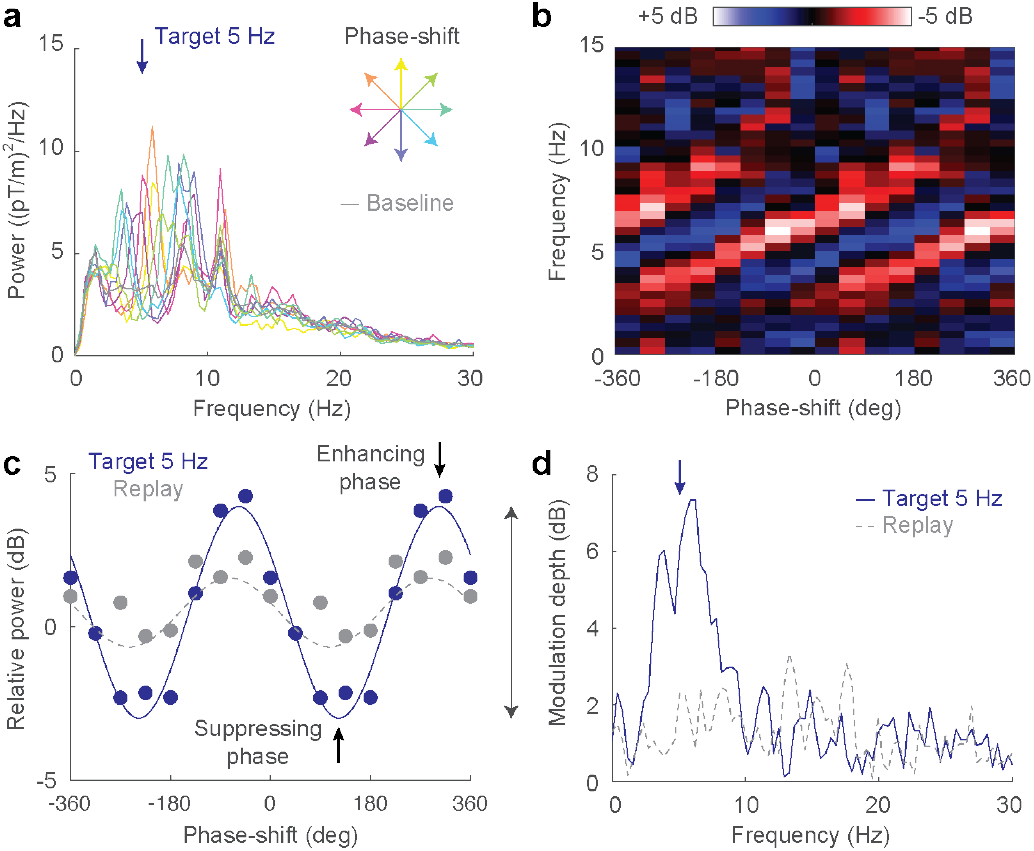
Brain-responsive music modulates neural oscillations. (a) Power spectrum of an MEG sensor over temporal cortex while a subject listens to brain-responsive music targeting 5 Hz with eight different phase-shifts. (b) Power modulation relative to baseline for different frequencies and phase-shifts. (c) A sinusoid fit of relative power was used to define modulation depth for each frequency. (d) Modulation depth was peaked around the target frequency. Open-loop replay of the same music a second time resulted in greatly diminished modulation of power.

The resulting spectra are characterised by peaks and troughs that depend on the phase-shift imposed within the feedback loop. These are revealed more clearly in a frequency-phase plot (Figure 3b) of power relative to the baseline condition (unmodulated music). Note that the frequencies where power was enhanced or suppressed became higher with increasing phase-advance in the feedback loop leading to a pattern of diagonal strips. This pattern is characteristic of delayed closed-loop feedback, with the slope of the stripes determined by the group delay in the feedback loop. At each frequency we calculated a peak-to-trough modulation depth from a sinusoidal fit of relative power against phase-shift (Figure 3c; blue line). Modulation depth was greatest around 5 Hz, the frequency targeted in this case by the closed-loop algorithm (Figure 3d; blue line). Note that a modulation depth of 7 dB corresponds to a 5x difference in power between enhancing and suppressing phases at the target frequency, representative of the typical effects observed in our MEG experiments.

Interestingly, such strong modulation of brain signal power was not seen when we replayed the same music a second time as an open-loop stimulus (Figure 3c,d; grey lines). Although our replay conditions comprised identical music to the corresponding closed-loop conditions, the timbral manipulations were no longer synchronized with endogenous brain activity so their effect on spectral power was attenuated. Note that the difference between closed-loop control and open-loop replay is particularly pronounced for suppressing phases in Figure 3c. This is to be expected since while it is possible to excite modes of a system with a feedforward control signal (e.g. entraining an oscillation with rhythmical stimulation), feedback control is generally required to suppress ongoing activity. The ability to systematically enhance or suppress activity at the target frequency represents a key advantage of closed-loop stimulation using brain-responsive music, which cannot be achieved with pre-recorded music that is not appropriately synchronized to ongoing brain rhythms.

### 3.2 Brain-responsive music can target different brain areas

To determine the spatial selectivity of power modulation that could be achieved with brain-responsive music, we examined the effect of synchronizing music to different MEG sensor locations. For these experiments, the timing of notes within piano music (Supp File 2) was manipulated based on the signal from a gradiometer located over either right temporal cortex or midline frontal cortex. We again targeted theta frequencies at 5 Hz and used six equally-spaced phase-shift conditions in eight subjects. We assessed the statistical significance of phase-dependent power modulation across the whole head in both sensor and source spaces relative to a shuffled surrogate distribution. Figure 4 shows the spatial distribution of significant phase-dependent power modulation when music was synchronized to right temporal cortex revealing a tight cluster in this area, as well as to a lesser extent in the corresponding contralateral location (likely due to interhemispheric synchronization of brain rhythms). By contrast, when the same music was synchronised to a midline frontal sensor, phase-dependent power modulation was instead seen in frontal but not temporal regions. This illustrates another key advantage of closed-loop stimulation, namely that a diffuse stimulus like music can nevertheless be targeted to specific brain areas by virtue of being timed to local activity sensed within that area.

**Figure 4.**
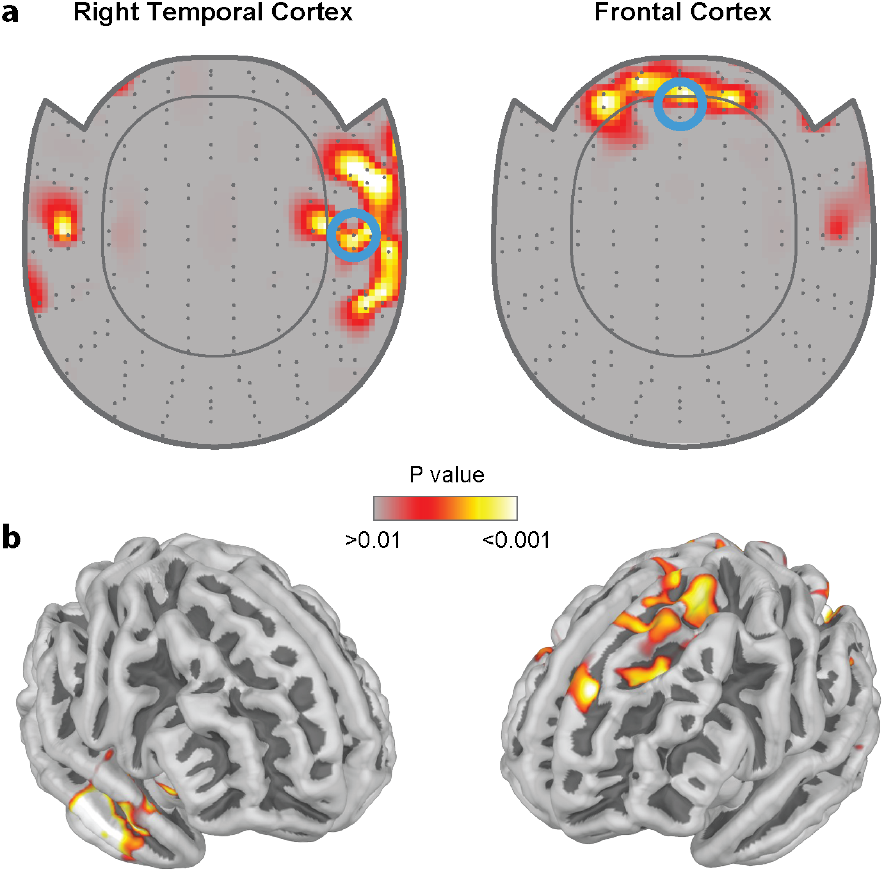
Brain-responsive music can target different brain areas. (a) Topographic maps of statistically significant phase-dependent power modulation resulting from music responding to sensor activity over right temporal cortex or frontal cortex (blue circles). (b) Equivalent statistical maps for activity projected into the source space. Data from 8 subjects.

### 3.3 Brain-responsive music can target different frequencies

To demonstrate that brain-responsive music could be targeted to different frequencies, we explored changing the centre frequency of the bandpass filter in the feedback loop from 5 Hz to 10 Hz. Figure 5a shows for an example subject how this could result in selective enhancement or suppression of the corresponding brain rhythm (in right temporal cortex). Figure 5b shows average phase-dependent power modulation for ten subjects, revealing that the effect of the music was greatest at the targeted frequency. Figure 5c shows the significant interaction between targeted frequency and power modulation at 5 Hz or 10 Hz (Two-factor ANOVA with replication, n = 10, F_1,36_ = 11.6, P = 0.002). For the corresponding replay conditions, no significant interaction was observed (F_1,36_ = 0.3, P = 0.6). These results demonstrate that brain-responsive music can be targeted to frequencies in either the theta or alpha band, although more research is needed to explore higher and lower frequencies.

**Figure 5.**
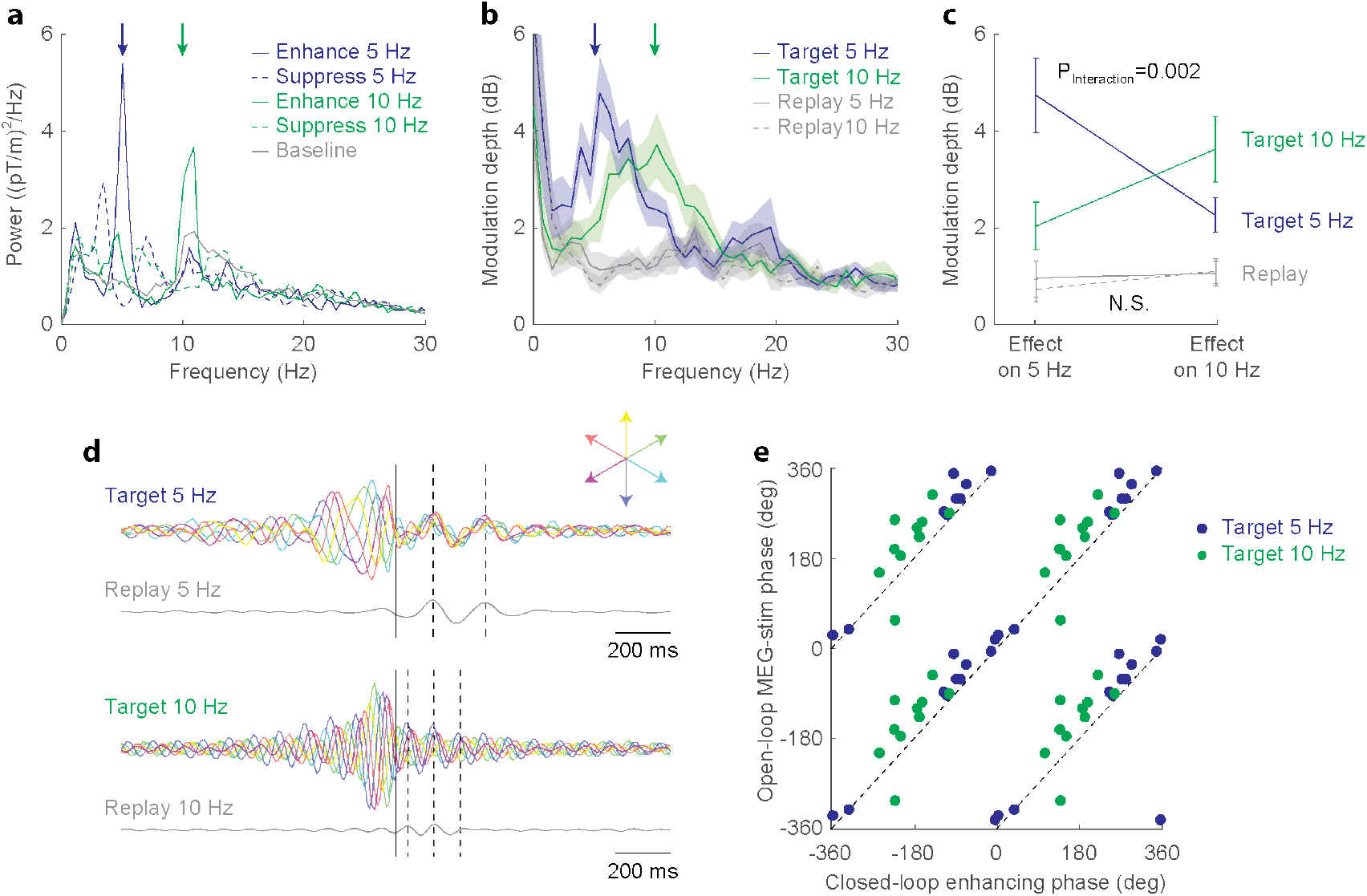
Brain-responsive music can target different frequency bands. (a) Example MEG power spectra for a subject listening to music that enhanced (solid lines) or suppressed (dashed lines) activity at 5 Hz (blue) or 10 Hz (green). (b) Average modulation depth for 10 subjects. (c) Phase-dependent modulation was significantly greater at the frequency (5 Hz or 10 Hz) that was targeted by the music. (d) Cross-correlations between brain signal and stimulus intensity under closed-loop conditions (coloured lines) revealed an entrainment that is predicted from the open-loop responses observed during the replay condition (grey lines). (e) Maximal closed-loop enhancement was seen with a filter phase advance that equalled the stimulus-brain phase delay as assessed from the open-loop data.

### 3.4 The optimal phase for closed-loop enhancement can be predicted from open-loop responses

Figure 5c shows, for an example subject, cross-correlations between the MEG signal and the output of the filter/phase-shift algorithm (which controlled the musical stimulus) for each of the six different phase conditions (*coloured lines*) with two target frequencies. Negative time-lags (left of time-zero) reveal the effect of preceding brain signals on the filter output and show, as expected, that the stimulus was maximal at different phases of the targeted rhythm. Positive time-lags (right of time-zero) reveal the effect of the stimulus on the subsequent brain signal and appear more clustered around a single phase, suggesting that the stimulus evokes or entrains a consistent neural response. The peaks of this response occur at the same time-lags as in the average cross-correlation for the open-loop replay conditions. Since the timing of the replay stimulus is unrelated to endogenous brain activity, these cross-correlations are similar to steady-state evoked potentials (albeit assessed using a slightly aperiodic stimulus). We speculated that the phase of this evoked potential might predict the closed-loop phase condition associated with maximal enhancement at the target frequency, since it effectively measures the total open-loop phase-delay between stimulus and resultant brain respose. Thus, an equal phase-advance imposed by our filter/phase-shift algorithm should result in a net closed-loop phase-shift of zero corresponding to a positive feedback.

To test this we plotted, for each subject and frequency condition, the phase-delay of the open-loop stimulus-evoked potential (calculated from the cross-spectrum between the replay brain signal and the stimulus) against the phase-advance associated with maximal power at the target frequency (calculated from a sinusoidal fit of spectral power for the different closed-loop phase-shift conditions). As shown in Figure 5d, these points are clustered around the line of equality. This result provides a simple method of calibrating, for a given individual and target frequency, the required phase-shift for maximal enhancement or suppression. By delivering a suitable open-loop stimulus around the target frequency (which can be achieved by providing a white noise input to our filter/phase-shift algorithm), we can assess the phase-delay of the evoked response and set the closed-loop phase-advance to this value (for enhancement) or this value plus 180° (for suppression). This calibration method was used for the subsequent EEG experiments described below.

### 3.5 EEG results

In order to translate brain-responsive music to a wearable system suitable for home-use, we tested whether EEG signals recorded using a reduced montage could be used to control brain-responsive music. For our first EEG experiments we used a wired system to record differential EEG between the forehead and right mastoid and targeted theta frequencies at 6 Hz. Figure 6a,b shows that brain-responsive music was able to produce an enhancement or suppression of ∼20% power at this frequency which, although smaller than the modulation we achieved in MEG experiments, was nevertheless highly significant from equivalent open-loop replay conditions (Two-tail paired t-test, n = 30, Enhance: t_29_ = 5.8, P = 2.7×10^−6^, Suppress: t_29_ = 5.8, P = 2.5×10^−6^).

**Figure 6.**
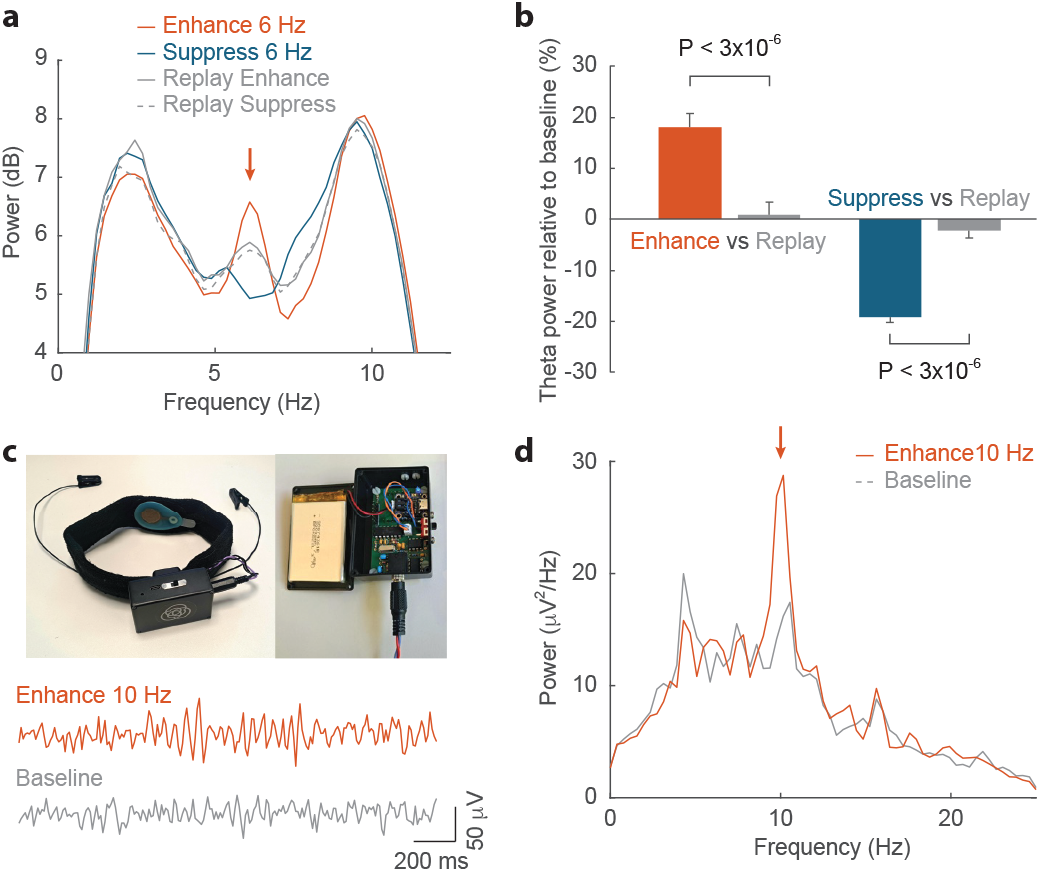
Brain-responsive music using EEG. (a) Average EEG spectra for 40 subjects listening to brain-responsive music targeting 6 Hz. (b) Significant enhancement and suppression was observed relative to open-loop replay of equivalent music. (c) Wearable EEG headband with dry electrode and wireless transmission of data. (d) Enhancement of alpha activity demonstrated using the wearable system.

Encouraged by these results, we have designed and built a battery-powered wireless device using a dry forehead EEG electrode and ear-clips for ground and reference (Figure 6c). Our device relays the EEG signal as Bluetooth MIDI messages which can be received directly by music software running on a computer or mobile device with low latency (∼10 ms). Figure 6d shows example power spectra for a subject listening to brain-responsive music designed to enhance alpha activity at 10 Hz. A video showing brain signal and spectral changes as the music alternates between enhance and baseline epochs is included as Supp File 3. This piece was composed by one of the authors (AJ) and incorporates multiple instrumental tracks to which a number of different brain-responsive timbral and timing manipulations are applied in real-time. While still crude in comparison to the sophistication of professional music, these results demonstrate the potential of brain-responsive music to have a pronounced impact on brain oscillations using a pleasant and unobtrusive stimulus.

## Discussion

The intersection of music and neurotechnology has a long and distinguished history. Ever since Adrian and Matthews (1934) found that the Berger rhythm could be ‘made audible by using a horn loud speaker’[19], neuroscientists and musicians have been fascinated by the concept of sonifying brain activity. Encephalophones[20], usually controlled by the bandpower of EEG rhythms, have been used as live performance instruments as well as for neurofeedback, while intracranial recordings have allowed a participant in the BrainGate2 trial to play music through a brain-controlled piano keyboard[21]. However, recent advances in both consumer BCI technologies and digital audio software may mean that their synergy is ready for mainstream applications. While volitional brain-control of musical instruments can find uses as an assistive device or performance novelty, we believe that modulating music with signals from the listener’s brain has further potential as an unobtrusive neuromodulation modality suitable for mass adoption. Building on principles derived from closed-loop neurostimulation and leveraging the capabilities of modern DAW software, we have shown how brain-responsive music can enhance or suppress oscillatory activity with spectral and spatial selectivity, without the need for active engagement from the user. Given that we listen to music on average for around 20 hours per week, this approach may allow neuromodulation therapies to be delivered for significant durations in home settings using a pleasant stimulus that does not interfere with users’s daily activities. Future work will explore whether such a therapy can provide equivalent or improved efficacy compared to conventional CLAS protocols, neurofeedback, open-loop sensory entrainment and other non-invasive brain stimulation methods that modulate neuronal oscillations for therapeutic applications in conditions ranging from ADHD[22], sleep[6], memory[5], pain relief[23] and more.

Importantly, the systematic effects of brain-responsive music on MEG and EEG power spectra were not observed when subjects listened to identical music as an open-loop stimulus. Brain-responsive music therefore appears to have effects on the brain that cannot be achieved with conventional recorded music, raising the intriguing possibility of future creative as well as therapeutic applications. A principal reason we listen to music is to regulate arousal and mood[24], brain states that are intimately tied to oscillatory processes. Indeed the extreme emotional and autonomic response of musical frisson or ‘chills’ is associated with a frontal theta signature[25]. It will therefore be interesting to explore whether brain-responsive musical elements can enhance and expand the expressive qualities of music by acting directly on brain oscillations associated with emotional states. Given the open-ended possibilities afforded by music as a stimulus space, determining how to integrate these elements in ways that are aesthetically effective will require as much musical as scientific experimentation. We hope that the tools and results described here can help provide both means and motive for future creative exploration at the crossroads of BCI and music technology.

## Acknowledgements

This work was supported by the EPSRC (EP/R511584/1). JR and AJ are directors and share-holders of Neudio Inc which is commercialising aspects of this technology.

## Supplementary Files

***Supp File 1***. *Example of brain-responsive timbral manipulation. This audio file contains first a period of baseline, unmodulated music, followed by timbral manipulation via a resonant low-pass filter with a cut-off frequency controlled by the brain signal*.

***Supp File 2***. *Example of brain-responsive timing manipulation. This audio file contains first a period of the original chord progression, followed by arpeggiation controlled by the brain signal*.

***Supp File 2***. *Example of brain-responsive multitrack music in which the timing and timbre of multiple instruments were controlled by the brain signal. This video shows the brain signal recorded using our wireless, wearable device, together power spectra and relative alpha power, compared for interleaved epochs of 10 Hz enhancement (red) and baseline (grey, music controlled by noise input)*.

